# PACAP alleviates the defective epididymis and sperm function in LPS-induced acute mice epididymitis

**DOI:** 10.1101/2025.02.23.639790

**Authors:** Zhang XueYing, Liu YaNan, Gong BenJiao, Wang XiaoXin, Liu XueXia, Liu FuJun

## Abstract

**Background:** To investigate the molecular mechanisms by which PACAP alleviates LPS-induced epididymitis, focusing on its anti-inflammatory and antioxidant pathways in the epididymal microenvironment.

**Methods:** A mouse model of acute epididymitis was induced by LPS injection. The study assessed the levels of inflammatory markers (IL-6 and TNF-α mRNA) in the cauda epididymis, sperm motility and morphology, morphological alterations in the epididymis, and the expression of inflammation and antioxidant-related genes. PACAP treatment was administered to evaluate its effects on these parameters.

**Results:** Results showed elevated levels of IL-6 and TNF-α mRNA in the cauda epididymis due to LPS injection. PACAP treatment effectively reduced these inflammatory markers and improved compromised sperm motility and morphology. PACAP also ameliorated morphological changes in the epididymis and mitigated the LPS-induced increase in leukocyte and macrophage markers. Gene expression analysis revealed that PACAP co-treatment suppressed LPS-induced upregulation of Il-6 and Tnf-*α* in caput tissues. In cauda tissues, PACAP significantly reduced the expression of LPS-elevated Tnf-*α* and IL-1β. PACAP up-regulated the antioxidant genes Cat and Sod1, which were down-regulated by LPS. IVF experiments demonstrated that PACAP restored the effects of LPS on sperm-egg fusion and embryo development. In conclusion: PACAP played a crucial role in maintaining epididymal function and sperm quality in the presence of inflammation.

## Introduction

Acute epididymitis was an inflammatory condition of the epididymis that can significantly impact sperm quality. The pathophysiology underlying this condition involved various mechanisms that led to alterations in sperm parameters, including concentration, motility, and morphology [1]. The inflammation associated with acute epididymitis can induce oxidative stress and the release of reactive oxygen species (ROS), which were detrimental to sperm function. Specifically, pathogens responsible for epididymitis can directly damage sperm quality through inflammatory mediators and cytokines, such as TNF-α and IL-6, which were known to disrupt spermatogenesis and sperm - 3 maturation processes [2].

The epididymis played a vital role in sperm maturation, and any disruption in its function can have long-term consequences for sperm quality [3]. Furthermore, the integrity of the epididymal epithelium was essential for proper sperm maturation. Damage to the principal cells of the epididymis, which secrete vital proteins and factors necessary for sperm maturation, can lead to impaired sperm motility and fertilization capacity [4]. In cases of obstructive azoospermia, which can be a consequence of acute epididymitis, sperm motility was notably affected, highlighting the importance of the epididymal environment in maintaining sperm functionality [5]. In addition to direct effects on sperm, acute epididymitis can also trigger an immune response that may further complicate recovery [6]. The presence of inflammatory cytokines can lead to the formation of anti- sperm antibodies, which can negatively impact sperm motility and overall fertility. Therefore, acute epididymitis has a significant effect on sperm quality and may cause serious fertility damage.

PACAP was an important neuroendocrine factor that was expressed at a higher level in the epididymis, suggesting that it may play an important role in the epididymal microenvironment for sperm maturation and in the process of sperm maturation [7]. Recent studies have highlighted the role of PACAP in enhancing fertility, particularly in the context of obesity-related reproductive dysfunction. For instance, Yan et al. demonstrated that PACAP treatment improved reproductive parameters in obese male mice, including testis morphology and sperm quality, through mechanisms involving the activation of Sirt1 and deacetylation of p53, which are critical for maintaining spermatogenesis and overall reproductive health **[8]**. Similarly, Shan et al. reported that PACAP mitigated oxidative stress and inflammatory responses in spermatogenic cells via the PAC1 receptor and the PKA/ERK/Nrf2 signaling pathway, further supporting its protective role in male reproductive health [9]. These findings suggested that PACAP not only played a role in sperm maturation but also in protecting the reproductive system from inflammatory damage, which maybe potential relevant in the context of acute epididymitis.

In this study, we administered PACAP to treat LPS-induced acute epididymitis and subsequently assessed tissue morphology, sperm functionality, molecular markers in the epididymis, as well as the expression of various inflammation molecules. Through a comprehensive analysis of phenotypes, we discovered that PACAP significantly mitigated the inflammation associated with LPS-induced epididymitis and alleviated damage to both epididymis and sperm function resulting from epididymitis. These findings provided crucial insights for future investigations into the role of PACAP in modulating epididymis function and its potential applications in anti-inflammatory therapies.

## Materials & Methods

### Animal experiments

The six-week-old male C57 mice (approximately 20–22 g) were procured from Beijing Vital River Laboratory Animal Technology company and housed in a specific pathogen-free (SPF) environment with a stable temperature of 22 (± 2)°C, humidity of 45 (± 5)%, and a light/dark cycle of 12 hours each. All mice were provided ad libitum access to food and water. All experiments conducted in this study received approval from the Medical Ethics Committee of Yantai Yuhuangding Hospital (2024-749).

Forty-eight mice were divided into four groups, with 12 mice per group (n=12 per group). The control group received an intraperitoneal injection of normal saline (0.9% NaCl). The PACAP group received an intraperitoneal injection of PACAP (HY-P0221, MedChemExpress LLC, Shanghai, China). The LPS-treatment group received an intraperitoneal injection of Lipopolysaccharides (LPS, HY-D1056, MedChemExpress LLC, Shanghai, China). The LPS+PACAP treatment group received both LPS and PACAP as administered above. In the preliminary experiment, four mice were randomly assigned to each group for sample collection to detect cytokine expression 6 hours after treatment in order to confirm the success of the mouse model. The remaining mice were treated for 72 hours and changes in parameters related to epididymis and sperm function were assessed. At the end, all experimental groups were euthanized using a sterile anesthetic solution of 1.25% 2,2,2-Tribromoethanol at a dosage of 0.2 ml/kg via intraperitoneal injection. Subsequently, one epididymis from each mouse was collected and fixed in Bouin’s solution (HT10132, Sigma, St. Louis, MO, USA) for immunochemistry staining and microscopic evaluation purposes; while the other epididymis was rapidly frozen in liquid nitrogen to facilitate subsequent mRNA and protein extraction. The epididymal spermatozoa were collected from the cauda epididymis and placed in PBS supplemented with 10% (w/v) bovine serum albumin (BSA). Several small incisions were made to release live sperm at 37 °C for 5 minutes. The sperm parameters were evaluated using Computer-assisted Sperm Analysis (CASA) system (Hamilton Thorne, Beverly, MA, USA).

### Morphological and indirect immunofluorescence (IF) analysis

Hematoxylin and eosin (HE) staining was performed according to our previous protocol for assessing morphological changes [10]. The embedded slides were de-paraffinized in three rounds of xylene and underwent a gradient series of ethanol washes from 100% to 90–75%. Subsequently, the slides were stained with hematoxylin for 10 minutes, followed by a 2-minute rinse in running water. Then, the slides were stained with eosin solution for 1 minute. After dehydration, the slides were examined using light microscopy (DM LB2, Leica, Nussloch, Germany). For immunofluorescence (IF) analysis, the de-paraffinized slides underwent antigen retrieval in citrate buffer solution (0.01 M, pH 6.0) using medium heat in a microwave for 20 minutes. Endogenous peroxidases were eliminated using 3% H_2_O_2_ (v/v), and non-specific protein sites were blocked with Tri-buffered saline (TBS) containing 3% (w/v) bovine serum albumin (BSA) at room temperature for one hour. The slides were then incubated overnight at 4 °C with primary antibody (Anti-AQP1, ab12504; Anti-KRT5, ab183336, Abcam, Cambridge, UK; Anti-CD45, DF6839; Anti-F4/80, DF2789, Affinity Biosciences, JiangSu, China). Following three rinses with TBS, the slides were incubated at 37 °C for one hour with FITC-labeled anti-rabbit IgG secondary antibody diluted at a ratio of 1:400. Nuclei staining was achieved by visualizing propidium iodide (PI; Invitrogen Carlsbad CA; concentration: 0.01 mg/ml). Images were captured using fluorescent microscopy equipment (Observer7 Carl Zeiss Jena Germany). Finally, the average numbers of positive-staining signals were counted and calculated using ImageJ software.

### Real-time quantitative PCR

The total RNA from each sample was extracted using the TRIzol reagent, following our previously published protocols (Zhang et al., 2023). Subsequently, cDNA synthesis was performed using the 5 × All-In-One RT MasterMix with AccuRT (G592, Abm, Jiangsu, China), according to the manufacturer’s instructions. PCR reactions were carried out with BlasTaq 2 × qPCR MasterMix (G891 Abm, Jiangsu, China) on an ABI Prism 7500 instrument (Thermo Scientific, Shanghai, China). The primer sequences are provided in Supplementary Table 1. *Actb* served as the internal control gene. The cycle threshold (CT) value for each gene was recorded. To determine gene expression within each group, ΔCT values were calculated by comparing the difference between CTgene and CTactb within the same group. For each gene analyzed, average CTactb values from the control group were used as reference. The results were presented as 2-^ΔCT^ and statistically analyzed using One-Way ANOVA.

### In vitro fertilization experiment

The female mice were induced to undergo superovulation by intraperitoneal injection of 5 IU pregnant mare serum gonadotrophin (PMSG) and human chorionic gonadotrophin (hCG) at 48- hour intervals. Cumulus oocyte complexes (COCs) were obtained after 14-17 hours following hCG injection. After euthanizing the female mice, the COCs were transferred to microscopic droplets of 200 μL HTF medium (72002, SUDGEN, Nanjing), which were then incubated at a temperature of 37°C with saturated humidity and in an atmosphere containing 5% CO_2_. Mouse epididymal sperm (2 ×10^5^ sperm) in each group were collected from the cauda epididymis and incubated in 200μl of C-TYH medium (72021, SUDGEN, Nanjing, China) at 37 °C in a 5% CO_2_ incubator for 0.5 h to complete capacitation. Subsequently, approximately 10^4^ sperm were transferred into pre-balanced HTF medium (72002, SUDGEN, Nanjing, China). After a 4-hour incubation period, fertilized oocytes were washed and transferred into KSOM medium (M1430, AIBEI, Nanjing, China). The average number of sperm binding to oocytes and the percentage of developed two-cell stage embryos were quantified.

### Statistical analysis

The data were presented as the mean ± standard deviation for triple replicates. All statistical analyses were conducted using GraphPad Prism 9 software. The mean values were determined using One Way ANOVA, and a p value less than 0.05 was considered statistically significant.

## Results

### 1. Characterization of the epididymis and sperm functionality in mice with acute epididymitis induced by LPS

The induction of acute epididymitis in mice was achieved by intraperitoneal injection of LPS. After a 6-hour period, there was a significant increase observed in the mRNA expressions of IL- 6 and TNF-a in the cauda epididymis of LPS-treated mice. Importantly, treatment with PACAP effectively attenuated the expression levels of *IL-6* and *Tnf-α* induced by LPS (Figure 1).

**Figure 1.**
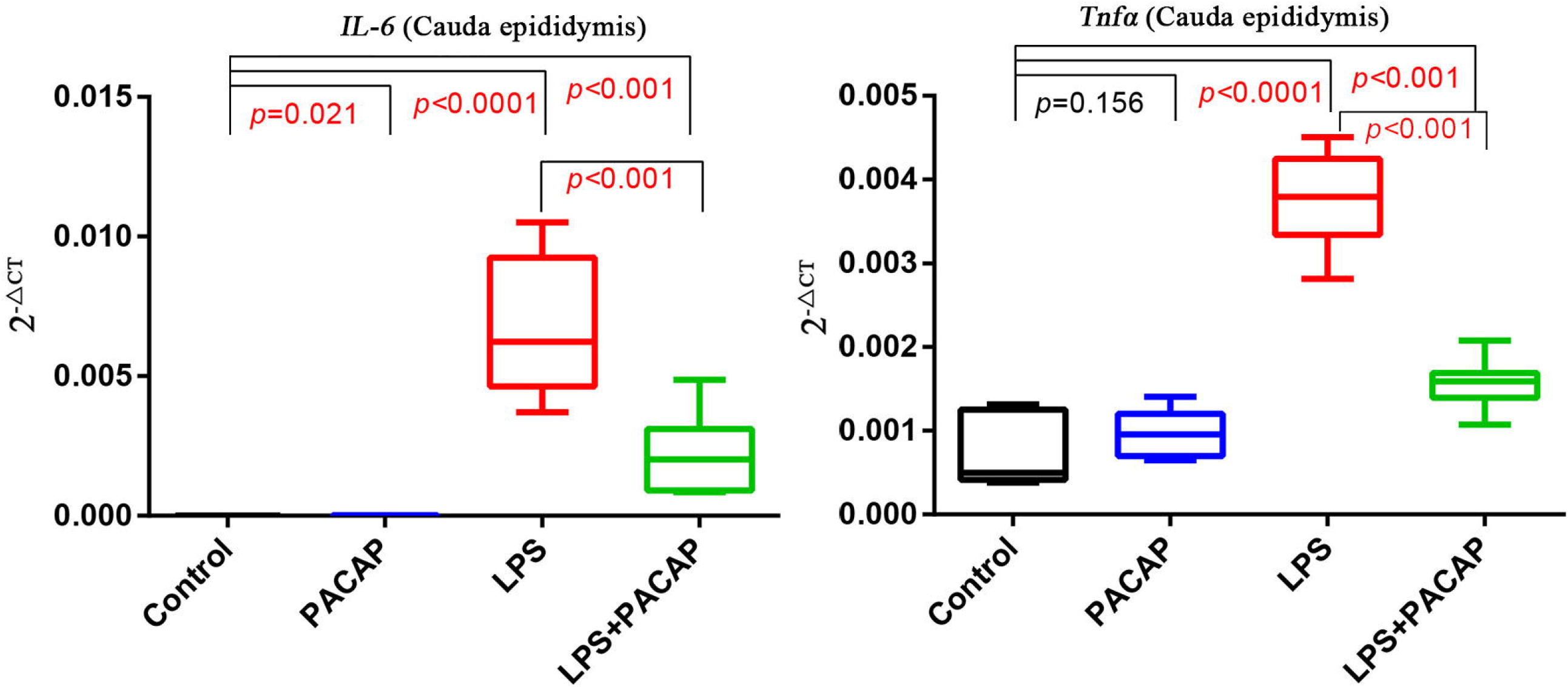
Gene expression of IL-6 and Tnf-*α* in caudal mice epididymis treated with LPS, PACAP or LPS+PACAP for 6 hours.

**Figure 2.**
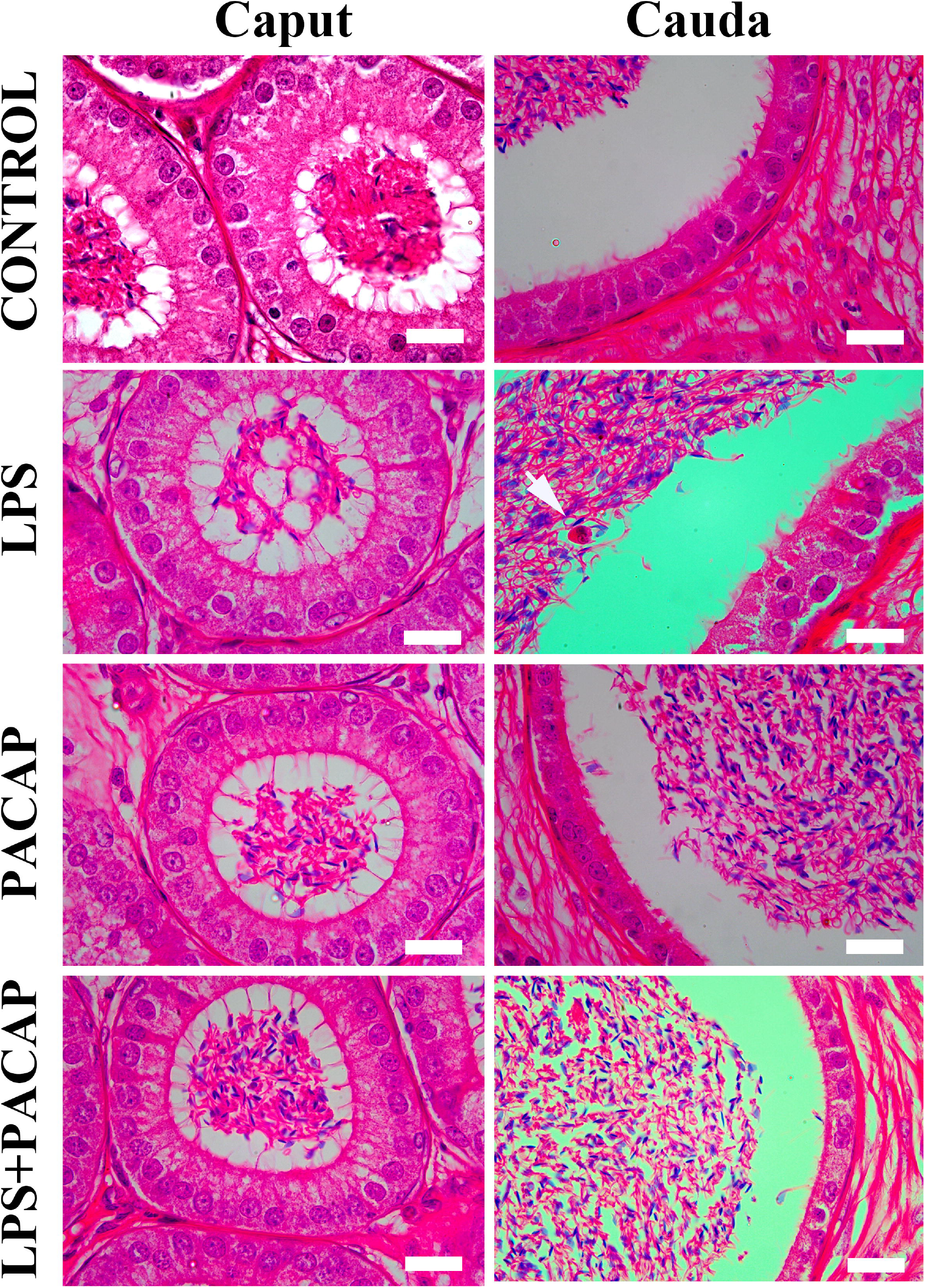
Morphological analysis of caput and cauda epididymis from different groups. Mice were administered LPS, PACAP, or a combination of LPS+PACAP for 72 hours. Samples of the cauda and caput epididymidis were then collected and stained with HE for histological observation. White arrows indicated inflammatory cells. Each bar represented 20μm

**Figure 3.**
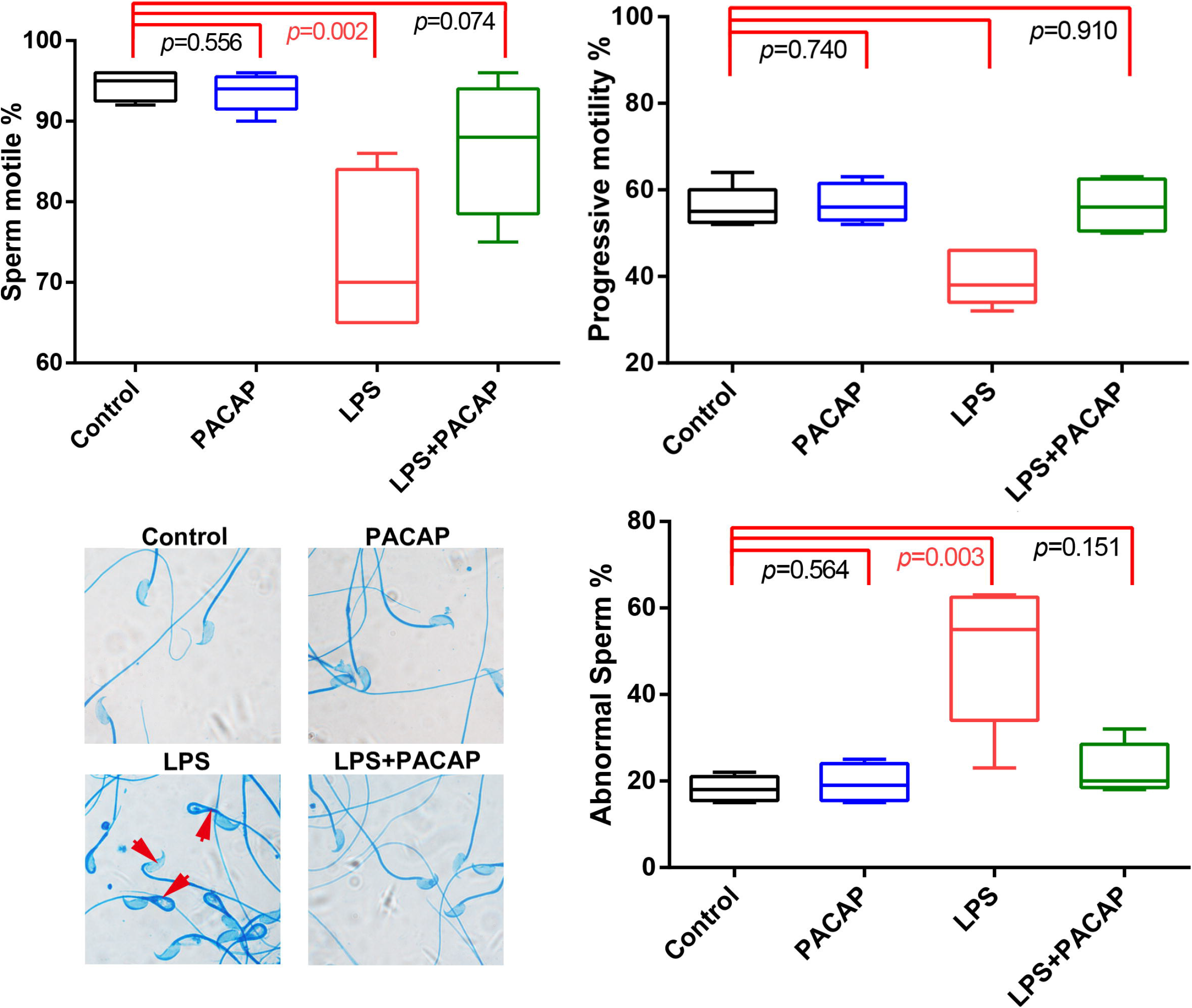
Characteristic of epididymal sperm viability, motility, and morphology in LPS and PACAP treated mice. The mice were treated with LPS or PACAP, and samples were collected 72 hours later. Data of each group were obtained from eight mice and analyzed by one-way ANOVA, p value less than 0.05 considered significance.

**Figure 4.**
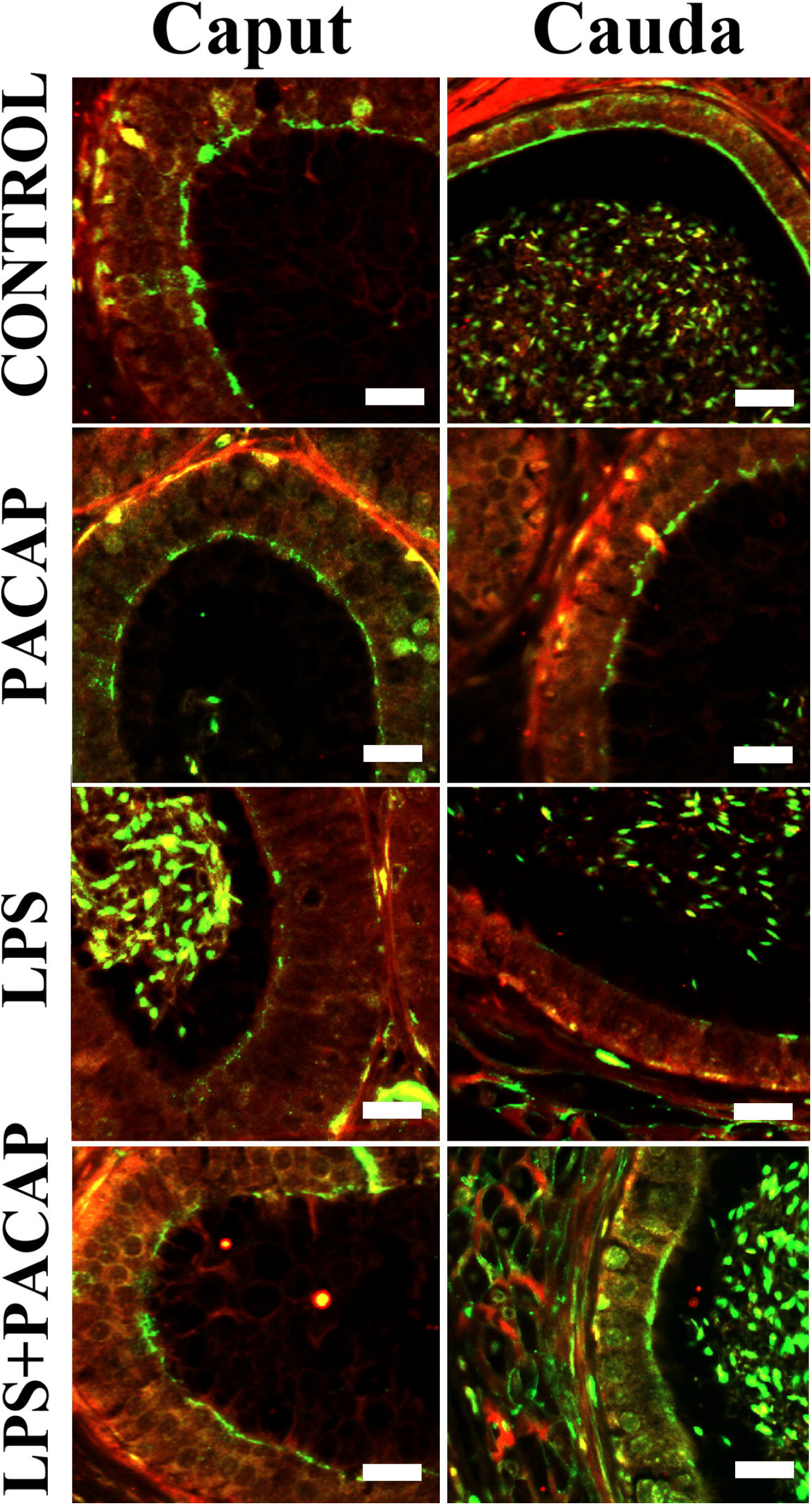
Expression of AQP1 in caput and cauda epididymis epithelium of LPS and PACAP treated mice. The mice were treated with LPS or PACAP, and samples were collected 72 hours later. The green signals indicated the AQP1 positive staining, and the red ones represented the PI stained nucleus. Each bar represented 10μm.

**Figure 5.**
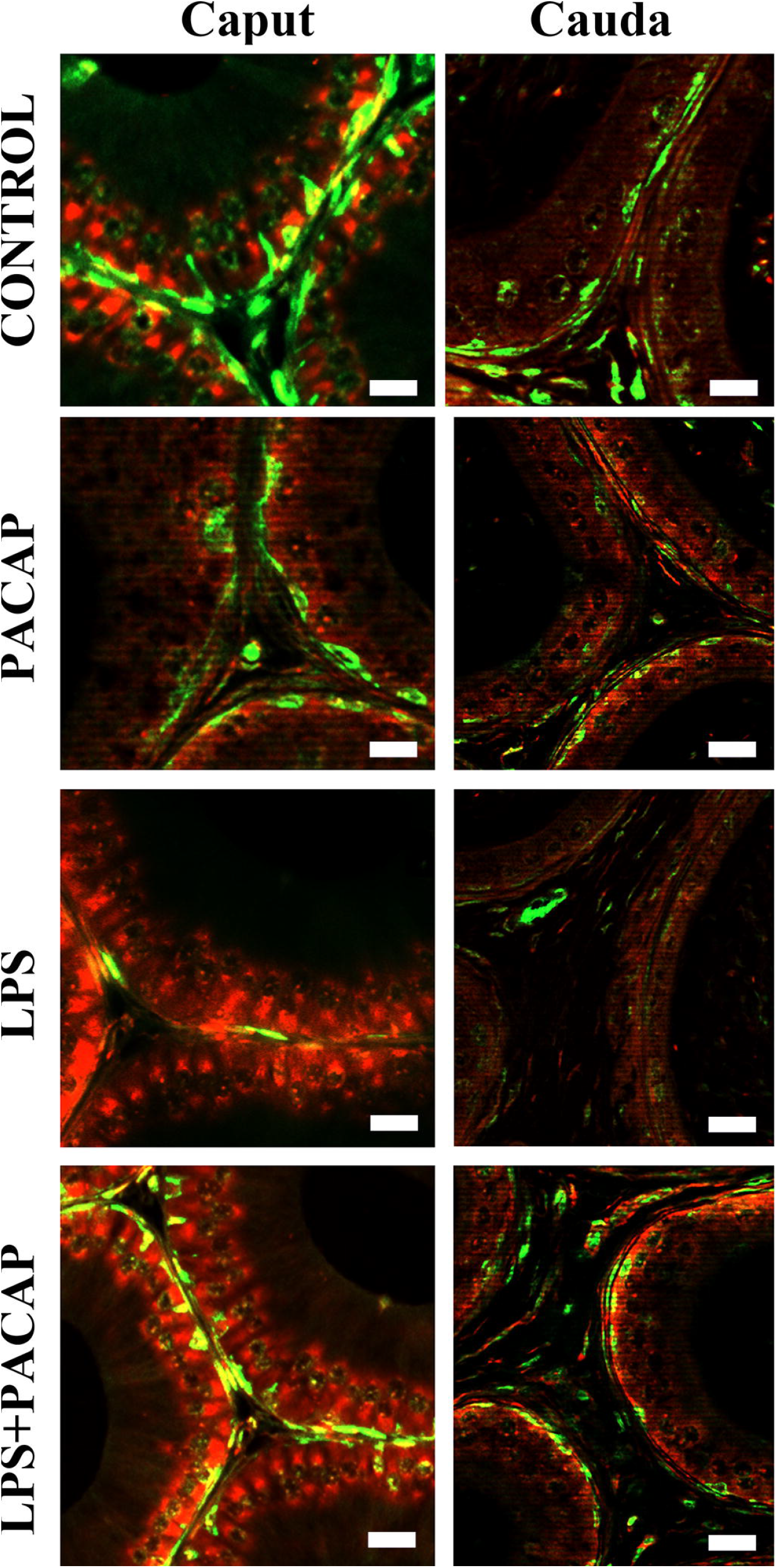
Expression of KRT5 in caput and cauda epididymis epithelium of LPS and PACAP treated mice. The mice were treated with LPS or PACAP, and samples were collected 72 hours later. The green signals indicated the KRT5 positive staining, and the red ones represented the PI stained nucleus. Each bar represented 10μm.

**Figure 6.**
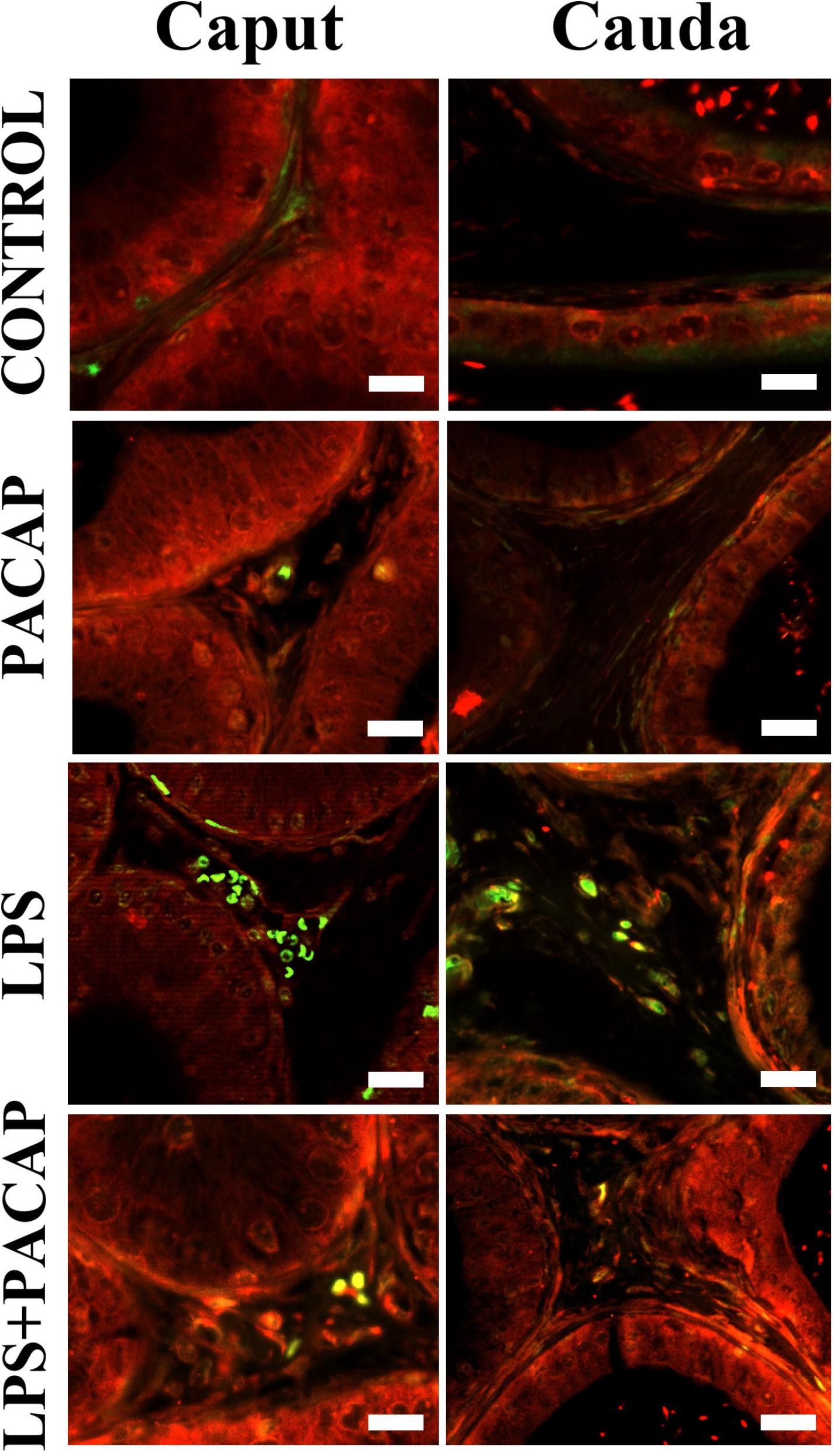
Expression of CD45 in epididymal caput and cauda epithelium under different treatments. CD45 was stained by FITC showing green fluorescence, and the nucleus was stained by PI. Each bar represented 10 μm.

**Figure 7.**
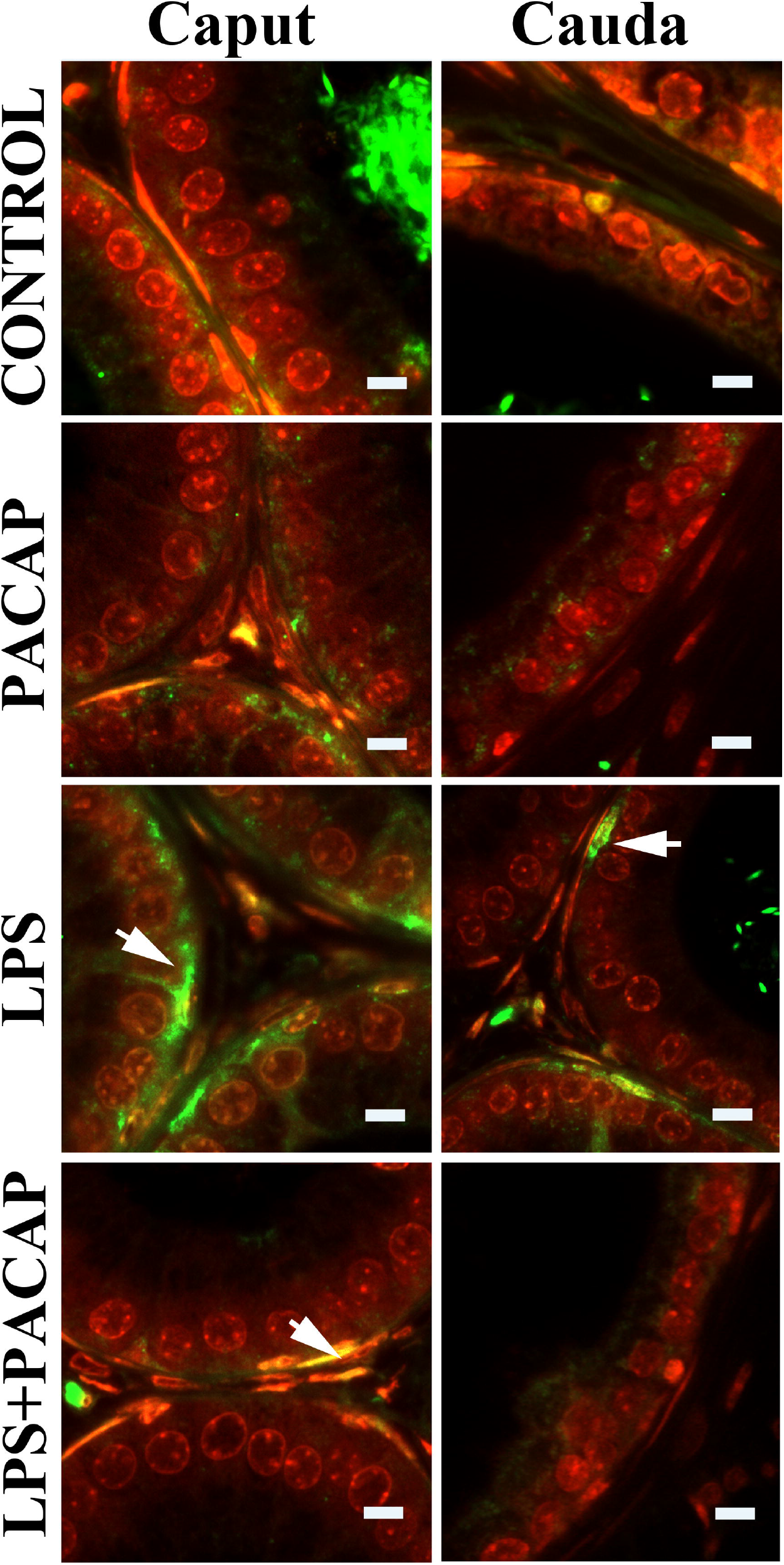
Expression of F4/80 in epididymal caput and cauda epithelium under different treatments. F4/80 was stained by FITC showing green fluorescence, and the nucleus was stained by PI. Each bar represented 10 μm.

**Figure 8.**
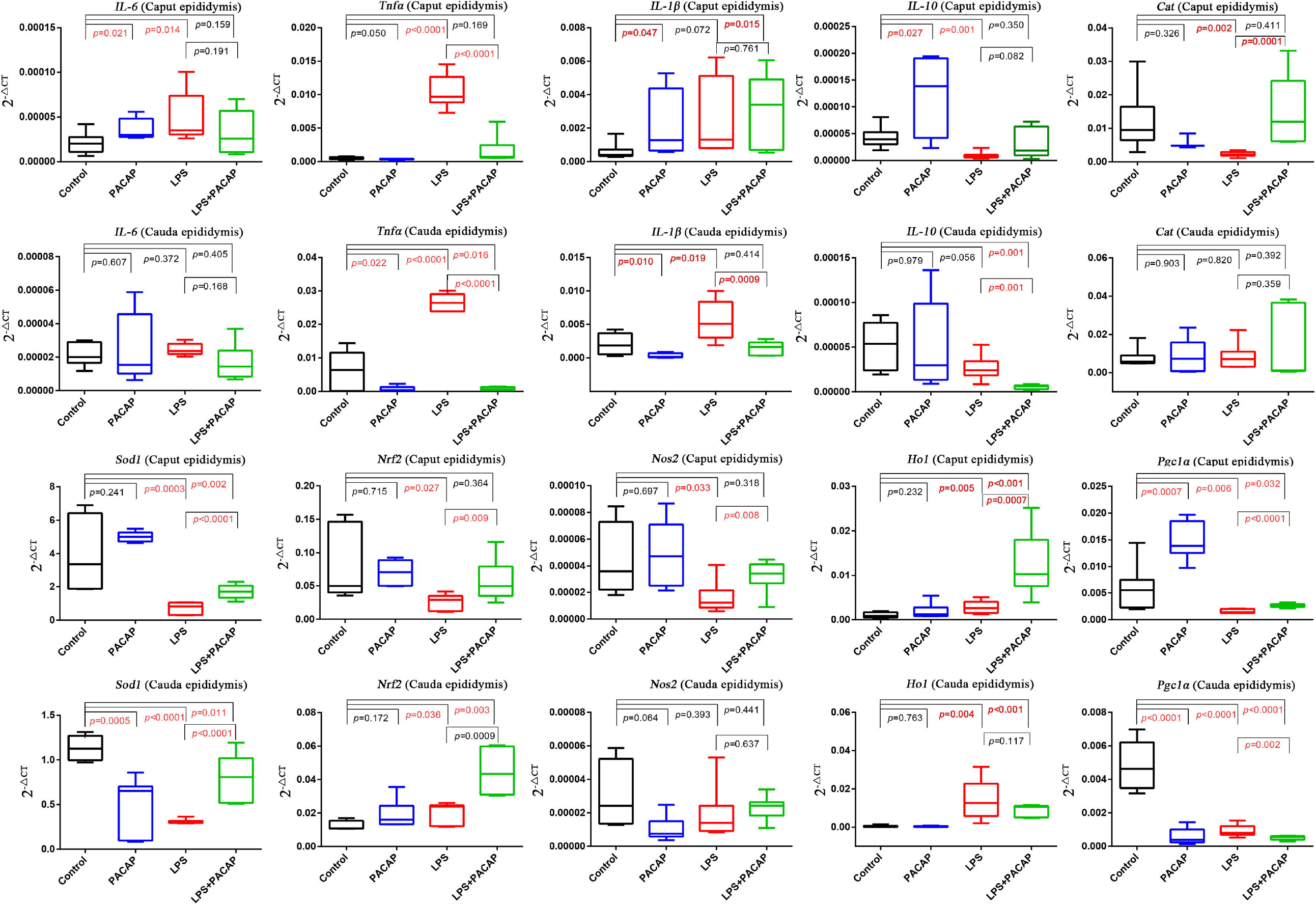
Expressions of key genes related to inflammation and antioxidant in the caput and cauda epididymis of different treatment groups.

**Figure 9.**
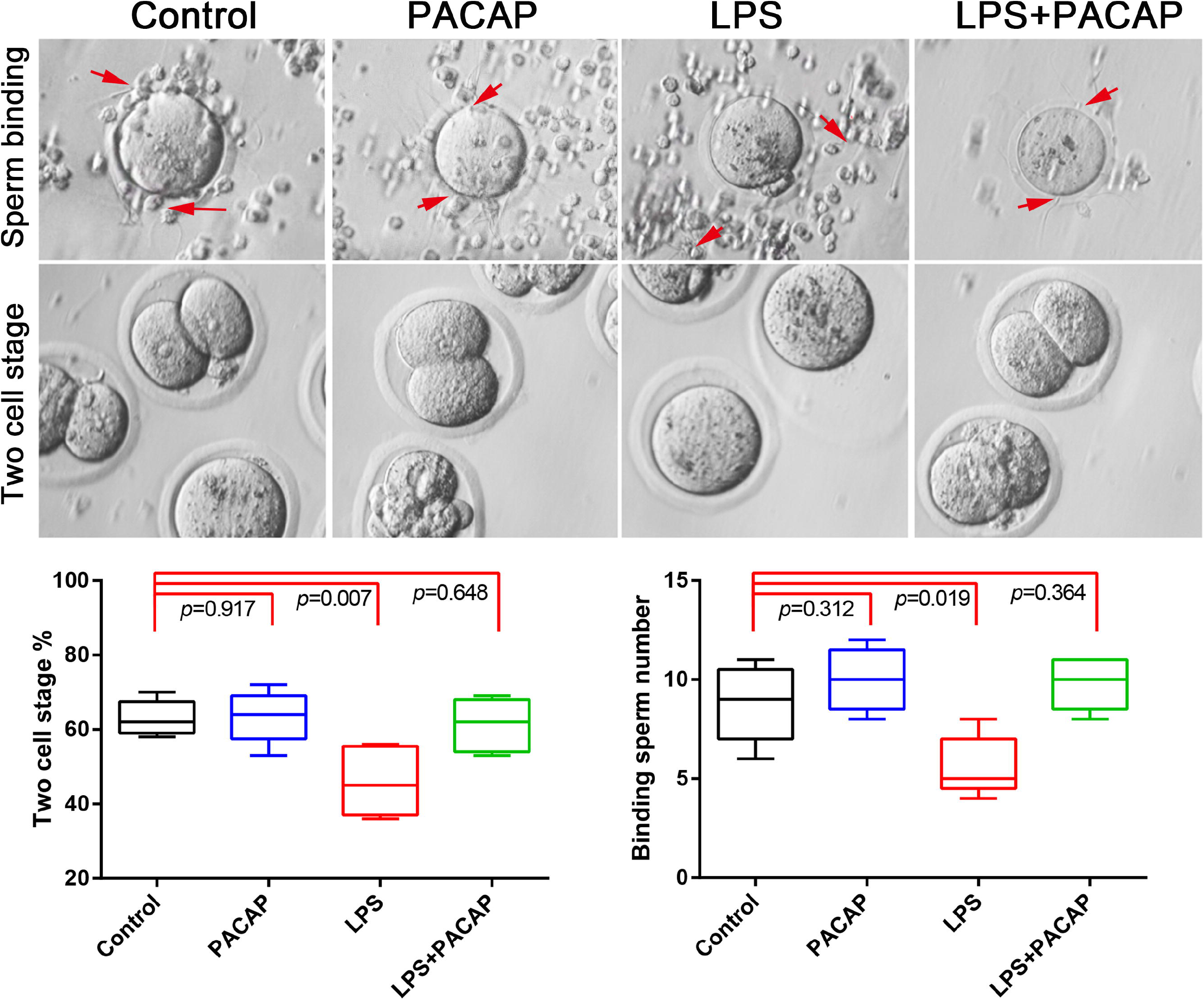
Epididymal sperm fertility analysis by in vitro fertilization experiments. Data were presented as the mean±SD of four mice in each group, and analyzed by one-way ANOVA, p value less than 0.05 considered significance. The arrows indicated the binding sperm.

The morphology of epididymal tissues was examined using HE staining. The results revealed that there were no significant alterations in the morphology of the caput and cauda epididymis following 6 hours of LPS treatment. However, after 72 hours of LPS treatment, scattered inflammatory cells were observed in the cauda epididymal lumen, and the epithelial cell layers were arranged disorderly and the stroma was proliferated. Notably, PACAP administration significantly ameliorated these alterations.

To obtain a more comprehensive understanding of the anti-inflammatory effects of PACAP in epididymitis, we opted to examine the epididymis and sperm samples obtained from mice subjected to LPS treatment for a duration of 72 hours. The testis/body weight ratio and sperm counts of mice in each group did not exhibit any significant alterations. However, the group treated with LPS demonstrated a notable reduction in both the percentage of motile sperm and progressive sperm motility, accompanied by an increase in abnormal bent sperm morphology. These parameters exhibited significant improvement following PACAP treatment.

### 2. Expressions of key proteins associated with epididymis function

The caput and cauda epididymis were subjected to immunofluorescence analysis, wherein the cell markers AQP1 and KRT5 were evaluated. AQP1 was predominantly expressed in principal cells of the caput and cauda epididymis within the stereocilia. Treatment with LPS significantly decreased the expression of AQP1 in both caput and cauda epididymis cells. KRT5 exhibited predominant expression in basal cells of the epididymis, which was also reduced by LPS treatment. Notably, there was a significant reduction observed in intracellular protrusions of basal cells as well. Following PACAP treatment, expressions of both AQP1 and KRT5 were significantly maintained compared to LPS treatment. These disparities suggested that LPS may impact the functionality of epididymal cells, consequently influencing the microenvironment for sperm maturation.

### 3. Expressions of CD45 and F4/80 in caput and cauda epdidiymis

Immunofluorescence labeling revealed the presence of leukocyte marker CD45 and macrophage marker F4/80 in both caput and cauda epididymis. LPS treatment induced a significant increase in CD45+ leukocytes within the interstitium of both regions, while only minimal levels were observed in the other three groups. F4/80+ macrophages were predominantly expressed on the basal side of epididymal epithelium, with their expression significantly elevated following LPS exposure. However, PACAP administration was able to effectively mitigate this increase.

### 4. Analysis of inflammation and antioxidant related genes in mice epididymis treated with LPS

The expression of inflammation and antioxidant-related genes in caput and cauda epididymis tissues from mice treated with PACAP or LPS for 72 hours was assessed using RT-PCR assays. In caput epididymis tissues, LPS treatment significantly increased the expressions of Il-6 and Tnf- *α*, while co-treatment with PACAP decreased their expressions, particularly resulting in a significant decrease in Tnf-*α* expression. IL-10 expression showed a significant reduction under LPS treatment but was up-regulated by co-treatment with PACAP. In cauda epididymis, Tnf-*α* and IL-1β exhibited significant up-regulation upon LPS treatment, which were significantly down- regulated by co-treatment with PACAP. The well-known antioxidant genes Cat and Sod1 were significantly down-regulated by LPS treatment in both caput and cauda epididymis tissues but were significantly up-regulated by PACAP treatment. Their upstream regulatory molecules Nrf2, Nos2, Ho-1, and Pgc1a displayed similar expression patterns in both caput and cauda epididymis.

### 5. PACAP reduced LPS-induced sperm function by in vitro fertilization experiments

The inflammation and oxidative stress induced by acute epididymitis can directly or indirectly impact sperm function, as evidenced by alterations in various sperm parameters. In order to assess this, we conducted in vitro fertilization (IVF) experiments using mice to evaluate sperm-egg binding and two-cell embryo development. Our findings revealed a significant reduction in the rate of both sperm-oocyte fusion and two-cell embryo formation upon exposure to LPS. However, no notable differences were observed between the control group and either the PACAP group or the LPS-PACAP group with regards to the number of sperm-egg fusions or the rate of two-cell formation.

## Discussion

The prevalence of acute epididymitis as a clinical condition affecting the male reproductive system can lead to various reproductive complications, including impaired epididymal function and altered sperm quality, ultimately impacting male fertility [11]. Due to its pleiotropic nature, PACAP, a widely distributed neuropeptide throughout the body, exerts anti-apoptotic, anti-inflammatory, and antioxidant effects [12]. The high expression of PACAP in the epididymis suggested its potential role in regulating epididymal function.

In this study, we successfully induced acute epididymitis in mice by intraperitoneally administering LPS. Our findings revealed a significant upregulation of *IL-6* and *TNF-α* gene expression levels in the cauda epididymis six hours post-LPS treatment, confirming the successful establishment of our experimental model. To comprehensively evaluate the impact of LPS treatment on epididymal function and sperm quality, our research focused on analyzing the expression levels of key molecules associated with epididymis function treated with LPS for a duration of 72 hours. Additionally, we assessed their influence on sperm performance and explored the protective role of PACAP. Following a 72-hour treatment of the epididymis with LPS, no notable decreases were detected in either testicular mass or sperm concentration. This outcome may be ascribed to the limited length of exposure. However, there was a significant impairment in the percentage of motile sperm and progressive motility sperm, leading to a notable increase in the proportion of abnormal spermatozoa. Treatment with PACAP did not elicit any significant changes; however, its combination with LPS exhibited a protective effect against LPS-induced damage to sperm. This suggested that PACAP may confer protection to sperm by modulating or safeguarding the microenvironment crucial for sperm maturation within the epididymis [5]. To elucidate the alterations in epididymal function during this process, we assessed the expression levels of AQP1 and KRT5 in the epididymis [13]. Our findings unveiled a significant reduction in AQP1 expression within both the caput and cauda epididymis following LPS treatment. This implied a compromised secretory function of epididymal epithelial cells, which may subsequently disrupt the essential microenvironment stability required for sperm maturation [14]. KRT5 served as a distinctive protein marker for basal cells in the epididymis, exhibiting clear expression on the basolateral side [15]. Following LPS treatment, there was a significant reduction in the number of KRT5-positive cells on the basolateral side, indicating a substantial impact of LPS on basal cells and potentially inducing apoptosis in epididymal cells. However, PACAP exerted a protective effect, as evidenced by no significant decrease in the expression levels of AQP1 and KRT5 when LPS and PACAP were combined.

Immunofluorescence assays revealed a significant upregulation of CD45 and F4/80 following LPS administration, indicating an enhanced inflammatory response [16]. In contrast, the administration of PACAP failed to induce such augmentation. RT-PCR experiments showcased a substantial elevation in pro-inflammatory cytokines including *IL-6, TNF-α*, and *IL-1β* upon LPS stimulation; meanwhile the antioxidant enzymes *Cat* and *Sod1* displayed a significant reduction. The protective effects of PACAP were exerted through diverse mechanisms involving the regulation of oxidative stress, a crucial factor contributing to cellular harm and impairment [17]. Studies have shown that PACAP can effectively mitigate cell death triggered by oxidative stress in different cell populations such as neurons and endothelial cells. For example, Douiri et al. demonstrated that PACAP exerted a protective effect on cultured rat astrocytes against oxidative damage induced by hydrogen peroxide, indicating its potential as a potent antioxidant [18]. Similarly, Horváth et al. reported that PACAP attenuated oxidative stress in renal and hepatic cells, highlighting its broad protective capabilities against oxidative insults [19]. This antioxidative action was crucial for epididymal cells, which were susceptible to oxidative stress due to their exposure to reactive oxygen species (ROS) during sperm maturation.

ACAP has been acknowledged for its potent antioxidant properties, which play a crucial role in safeguarding cells against oxidative damage. The ability of PACAP to augment the activity of endogenous antioxidant systems, such as superoxide dismutase and catalase, further emphasized its protective function against oxidative stress. In the context of the epididymis, where spermatozoa were particularly susceptible to oxidative damage due to their high content of polyunsaturated fatty acids, the anti-oxidative effects of PACAP could be indispensable for maintaining sperm viability and functionality [19].

The protective effect of PACAP was also evident in the enhancement of sperm quality observed in the IVF experiment. LPS significantly reduced sperm-egg binding and impaired two-cell development, indicating that LPS-induced inflammation influenced the expression of crucial molecules involved in sperm fertility. This finding also provided valuable insights for further comprehensive investigation into key molecules associated with sperm quality. The protective effect of PACAP on epididymitis was confirmed in this study through the detection of various molecules associated with epididymal function in the epididymitis model, providing crucial insights for further research on the anti-inflammatory properties of PACAP and its role in male reproduction, particularly in regulating sperm function.

## Conclusions

In summary, our study highlighted the protective role of PACAP in LPS-induced acute epididymitis. PACAP emerged as a potential therapeutic agent capable of mitigating the detrimental effects of inflammation on epididymal function and sperm quality. This finding underscored the importance of neuropeptides in maintaining male reproductive health and suggested future avenues for exploring PACAP-based therapies to treat epididymitis and associated fertility issues.

## Supporting information

Supplementary Table 1. Lists of primers for genes related to inflammation and antioxidant

## Acknowledgements

Not applicable.

