## Supplementary Table 1. Lists of primers for genes related to inflammation and antioxidant for "PACAP alleviates the defective epididymis and sperm function in LPS-induced acute mice epididymitis"

| **Gene name** | **Primer** |
|---|---|
| mActb-F | GCAGCTCAGTAACAGTCCGC |
| mActb-R | AGTGTGACGTTGACATCCGT |
| qmNrf2-F | TAGATGACCATGAGTCGCTTGC |
| qmNrf2-R | GCCAAACTTGCTCCATGTCC |
| qmHO-1-F | GATAGAGCGCAACAAGCAGAA |
| qmHO-1-R | CAGTGAGGCCCATACCAGAAG |
| qmINOS-F | GTTCTCAGCCCAACAATACAAGA |
| qmINOS-R | GTGGACGGGTCGATGTCAC |
| qmSOD1-F | AACCAGTTGTGTTGTCAGGAC |
| qmSOD1-R | CCACCATGTTTCTTAGAGTGAGG |
| qmCAT-F | TGGCACACTTTGACAGAGAGC |
| qmCAT-R | CCTTTGCCTTGGAGTATCTGG |
| qmPGC1α-F | TATGGAGTGACATAGAGTGTGCT |
| qmPGC1α-R | GTCGCTACACCACTTCAATCC |
| qmIL-1β-F | GAGTGTGGATCCCAAGCAATA |
| qmIL-1β-R | TCCTGACCACTGTTGTTTCC |
| qmIL-10-F | CCCAGAAATCAAGGAGCATTTG |
| qmIL-10-R | CACCTTGGTCTTGGAGCTTAT |
| qmTnfa-F | CATCTTCTCAAAATTCGAGTGACAA |
| qmTnfa-R | TGGGAGTAGACAAGGTACAACCC |
| qmIL-6-F | GAGGATACCACTCCCAACAGACC |
| qmIL-6-R | AAGTGCATCATCGTTGTTCATACA |
